# Coordination of Alternative Splicing and Alternative Polyadenylation revealed by Targeted Long-Read Sequencing

**DOI:** 10.1101/2023.03.23.533999

**Authors:** Zhiping Zhang, Bongmin Bae, Winston H. Cuddleston, Pedro Miura

## Abstract

Nervous system development is associated with extensive regulation of alternative splicing (AS) and alternative polyadenylation (APA). AS and APA have been extensively studied in isolation, but little is known about how these processes are coordinated. Here, the coordination of cassette exon (CE) splicing and APA in *Drosophila* was investigated using a targeted long-read sequencing approach we call Pull-a-Long-Seq (PL-Seq). This cost-effective method uses cDNA pulldown and Nanopore sequencing combined with an analysis pipeline to resolve the connectivity of alternative exons to alternative 3’ ends. Using PL-Seq, we identified genes that exhibit significant differences in CE splicing depending on connectivity to short versus long 3’UTRs. Genomic long 3’UTR deletion was found to alter upstream CE splicing in short 3’UTR isoforms and ELAV loss differentially affected CE splicing depending on connectivity to alternative 3’UTRs. This work highlights the importance of considering connectivity to alternative 3’UTRs when monitoring AS events.

## Introduction

mRNA transcripts are subject to a variety of co/post-transcriptional processing events in metazoan cells including alternative splicing (AS) and alternative polyadenylation (APA). These events are highly regulated during development and cell differentiation, including during neuronal differentiation^1, 2^. More than 70% of protein-coding genes in mammals and about half in flies harbor more than one functional polyadenylation site (polyA site)^3–5^. APA occurring in the terminal exon that alters the length of the 3’ untranslated region (3’UTR) is called tandem 3’UTR APA^6^. Transcripts of APA regulated genes cleaved at the distal polyA sites are highly enriched in the nervous system^5, 7, 8^. In *Drosophila*, the neuron-specific RNA binding protein Embryonic Lethal Abnormal Visual System (ELAV) is the major determinant of long 3’UTR expression in neurons via APA regulation^9–13^. In mammals, analogous roles in neural-specific 3’UTR lengthening has been proposed for Hu proteins and PCF11^14, 15^. Neural enriched long or extended 3’UTR mRNA isoforms have greater potential for regulation by RNA binding proteins (RBPs) and microRNAs. Long 3’UTR isoform-specific functions in the nervous system include dendrite pruning, axon outgrowth, reproductive behavior, and neural plasticity^16–21^. Disruption of APA regulators is linked to human disease, in particular members of the Cleavage Factor I (CFI) complex^22–24^. Mutations that cause the loss or gain of poly(A) sites also underlie several human disorders^25^. Genetic disruptions in 3’UTRs that affect APA are risk factors for brain disorders, such as *SNCA* in Parkinson’s disease^26, 27^.

In addition to its role in regulating APA, ELAV also widely regulates alternative splicing in the nervous system^10, 13^. A correlation between APA and AS has been reported in different model organisms^28, 29^. Many RBPs that were known to regulate AS have been found to be involved in regulation of APA^30, 31^. Cleavage and Polyadenylation factors have also been reported to bind coding regions to facilitate splicing^32^. The study of how AS and APA events are connected requires a sensitive sequencing method that can provide sequenced reads long enough to span from upstream of an alternative exon to the end of long 3’UTRs.

The last two decades have witnessed a rapid progress in next generation sequencing technologies that typically provide short-read sequencing information^33^. In contrast to the widely employed short read high-throughput sequencing technologies that generate reads < 150 nt in length, long-read sequencing techniques enable sequencing of full-length transcripts up to dozens of kilobases (kb)^34^. Currently, the two main platforms are Pacific Biosciences (PacBio) single-molecule real-time (SMRT) and Oxford Nanopore Technologies (ONT) nanopore sequencing platforms^35–38^. Long read sequencing holds great potential for understanding how different types of co/post-transcriptional processes are orchestrated across developmental stages, tissues, and cell types. Nascent RNAs have been examined using long-read sequencing technology to investigate how various RNA processing events are coupled^39^ and it has been found that upstream splicing events can alter downstream 3’ end processing in mammalian cells^36^.

A major drawback to transcriptome-wide long read RNA-sequencing is that full-length read information is mostly restricted to high abundance and relatively short transcripts. Enrichment strategies prior to long read sequencing permit sufficient depth of coverage for a targeted subset of genes. Several groups have successfully performed large-scale probe-based cDNA capture followed by PacBio long-read sequencing^40–42^. Others have employed cDNA capture coupled to long-read sequencing for smaller sets of genes on the nanopore platform^43, 44^. These studies have generally focused on new isoform discovery and exon connectivity, in particular alternative splicing. Fewer studies have employed these long-read approaches to quantify how alternative exons connect to alternative 3’UTRs regulated by APA^28^. This is a technically challenging problem given the especially long length of many 3’UTRs and the need to distinguish between tandem APA 3’UTRs that share a common region.

Our previous work showed that for the *Dscam1* gene in *Drosophila*, the alternative splicing of an upstream cassette exon (CE) was strongly influenced by whether it was connected to a long or short 3’UTR^18^. The neuron-specific RNA binding protein Embryonic Lethal Abnormal Visual System (ELAV) is a regulator of both AS and APA for many genes, including *Dscam1*^9–13^. *Dscam1* genomic long 3’UTR deletion and minigene reporter analysis showed that regulation of *Dscam1* CE splicing by ELAV required the presence of the long 3’UTR mRNA isoform.

Here, we set out to identify coordination of CE splicing and alternative 3’ end processing during *Drosophila* embryonic development. To accomplish this, we developed a targeted Nanopore long-read sequencing approach we call Pull-a-Long-Seq (PL-Seq). This approach facilitated the study of AS and APA coordination in a cost-effective and efficient manner. We used PL-Seq to identify 23 genes that exhibit 3’UTR connected AS in neuron-enriched tissues and quantify how ELAV regulates coordinated AS-APA. We also examined the cross-talk between AS and APA by genomic alteration of these events in *Drosophila* and quantifying their impact on one another.

## Results

### 3’UTR lengthening is significantly associated with CE regulation during embryonic development

Our previous work uncovered that coordinated AS and APA occurs during embryonic development for the *Dscam1* gene^18^. Browsing embryonic development short read RNA-Seq tracks (PRJNA75285)^45^ we identified another gene, *Khc-73*, that also shows coordinated upstream CE alternative splicing and 3’UTR lengthening (Fig. 1a). In this case, later developmental stages show increased inclusion of two upstream CEs (exons 12 and 15) that coincided with expression of the long 3’UTR isoform. We set out to identify more genes that undergo coordinated AS and APA in *Drosophila*. QAPA, a tool that enables estimation of alternative polyA site usage (PAU) of tandem APA events^46^, was used to quantify distal polyA site usage (dPAU) throughout embryonic development and in several dissected tissues. The distribution of dPAU across developmental stages and tissues revealed many genes shifting to distal polyA site usage later in development and in the nervous system, which is consistent with previous published observations (Fig. 1b)^5^. We compared the usage of distal 3’UTRs before (2-4 hr embryos) and after (16-18 hr embryos) establishment of the nervous system. Among 1951 expressed genes with multiple APA isoforms, 252 genes exhibited greater expression of the most distal 3’UTR in the later stage (lengthening), whereas 42 showed the opposite trend (shortening) (Fold Change>2 & *p*<0.05) (Fig. 1c).

**Figure 1.**
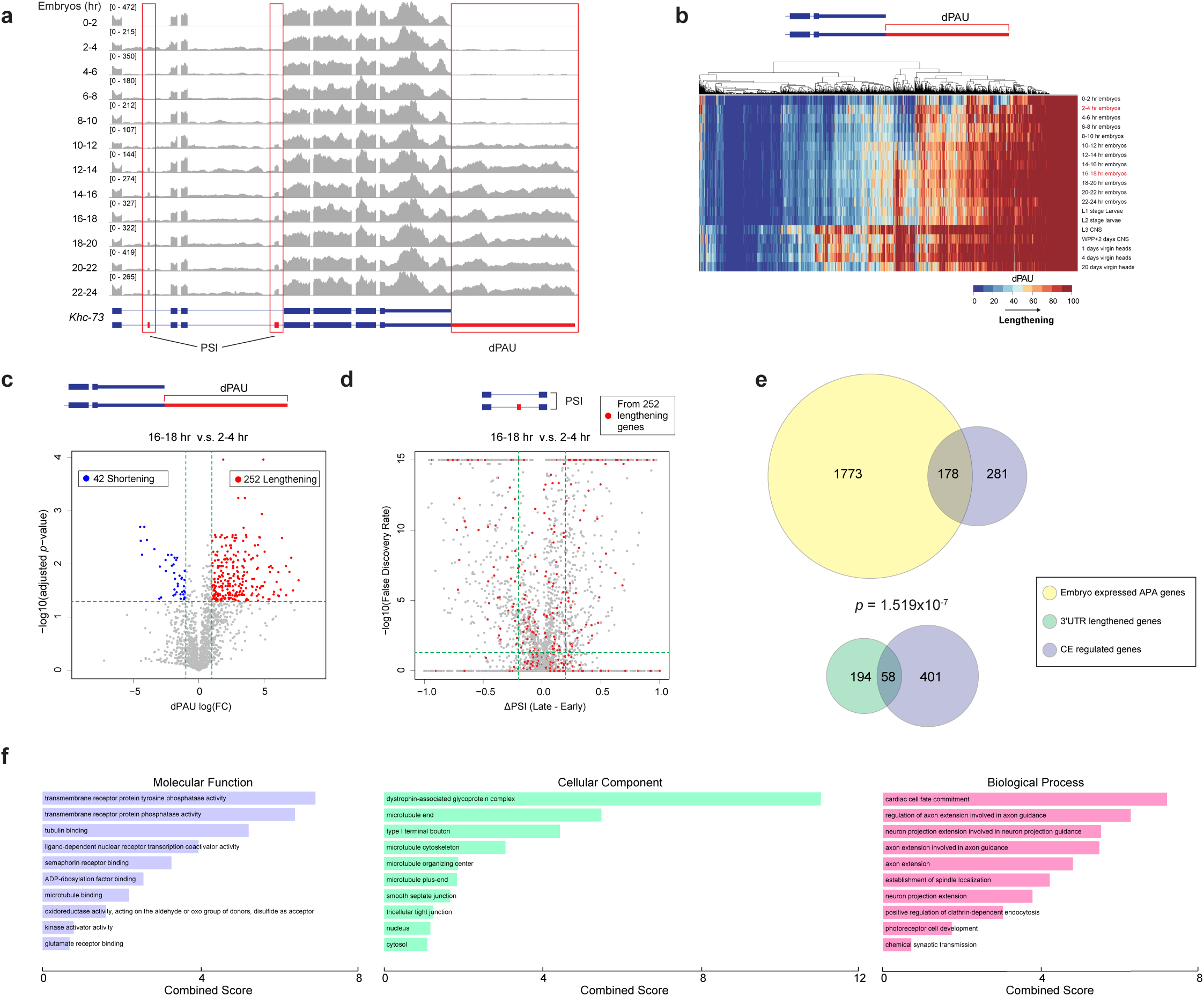
APA and AS analysis of short-read RNA-Seq data across development and tissues in *Drosophila*. (**a**) Short-read RNA-Seq data shows coordinated inclusion of CEs with 3’UTR lengthening during embryonic development for the *Khc-73* gene. (**b**) Distribution of Distal PolyA Site Usage (dPAU) values from different embryonic stages and larval/adult tissues. The value of dPAU falls into a range from 0 (no distal polyA site usage) to 100 (exclusive distal polyA site usage). (**c**) dPAUs compared between late-stage embryos (16-18 hr) and early-stage embryos (2-4 hr). *p* values are calculated from two-tailed student’s t-test, and then adjusted using FDR. Blue represents FDR adjusted *p*<0.05 and FC<0.5 while red represents adjusted *p*<0.05 and FC>2. FC, fold change. (**d**) Change in PSI compared between late-stage embryos (16-18 hr) and early-stage embryos (2-4 hr) for CEs. PSI ranges from 0 to 1, and ΔPSI is calculated as PSI (16-18 hr) – PSI (2-4 hr). Red dots represent CE events from the 252 3’UTR lengthening genes identified in (**c**). Horizontal dash lines in (c) indicate adjusted *p*=0.05 and in (d) indicate adjusted FDR=0.05. Vertical dash lines in (c) indicate FC=0.5 (left) and FC=2 (right) and in (d) indicate ΔPSI=-0.2 (left) and ΔPSI =0.2 (right). (**e**) Fisher’s exact test showing 3’UTR lengthening genes are significantly associated with regulated CE events in late vs early-stage embryos. (**f**) Gene Ontology analysis of the 58 genes exhibiting 3’UTR lengthening and regulated CE during embryonic development. Also see supplementary Fig. 1, 2.

Alternative splicing in the *Drosophila* central nervous system is widespread^47^. Using rMATS^48^, we detected a variety of regulated alternative splicing between 16-18 hr and 2-4 hr embryos. CE splicing was the largest group of AS events, with 358 exon skipping and 458 exon inclusion events affecting a total 446 genes (|ΔPSI|>0.2 and FDR<0.05) (Fig. 1d). For the 252 genes exhibiting 3’UTR lengthening during embryonic development, 58 also harbored one or more differentially regulated CEs (Fig. 1e). These 58 genes showed no preference for increased inclusion or skipping in the later developmental stage (58 CEs increased skipping, 82 CEs increased inclusion, *p*=0.6448, Fisher’s exact test). When compared with all the genes subject to APA in embryos, Fisher’s exact test showed that these 3’UTR lengthening genes were significantly associated with concurrently regulated CE splicing events (AS-APA) in the same host gene (Fig. 1e, *p*=1.519e-07). The CE splicing and APA developmental regulation trends for two of these genes, *Khc-73* and *Dys*, was confirmed using RT-PCR (Supplementary Fig. 1). Gene Ontology analysis revealed multiple categories of enrichment for the 58 genes, including molecular functions of phosphatase activity and receptor binding and biological processes of axon guidance (Fig. 1f).

For many APA regulated genes, isoforms using the distal polyA site are not highly expressed until adulthood in the brain (Fig. 1b). Thus, we performed an additional pair of comparisons between adult head and ovary samples, since ovary has been shown to generally lack long 3’UTR expression^5^. When compared with ovaries, 290 genes were found to have increased expression of the long 3’UTR isoform in heads, whereas only 8 showed the opposite trend (Supplementary Fig. 2a). Among these 290 3’UTR lengthening genes, 62 were found to also harbor one or more differentially regulated CE in heads compared to ovaries (Supplementary Fig. 2b). A significant association between 3’UTR lengthening and differentially spliced CEs was also revealed by Fisher’s exact test (Supplementary Fig. 2c, *p*=0.0002).

### PL-Seq reveals pairing of polyA site choice with CE alternative splicing for 23 genes

We wanted to understand how AS changes detected for the APA regulated genes are distributed between different 3’UTR isoforms. One might expect that if a given gene undergoes both AS and APA during embryonic development, the regulated CE could be 3’UTR independent or show a biased incorporation depending on the 3’UTR isoform. We performed an RT-PCR based nanopore sequencing approach targeted specifically for the *Khc-73* gene, which we previously developed for *Dscam1*^18^. We generated RT-PCR amplicons representing all 3’UTR isoforms that also capture the alternative splicing events for exons 12 and 15 (Uni), as well as amplicons that represent those associated with the extended 3’UTR sequence (Long). Nanopore sequencing revealed a significant difference in the PSI for both exons 12 and 15, with greater exon inclusion observed in the Long vs Uni amplicons (Supplementary Fig. 3a, b). This suggests that AS of these exons is connected to 3’UTR choice. Caveats of this approach include that it does not reveal the CE alternative splicing pattern specifically in short 3’UTR isoforms and the PCR amplification is performed for only a single gene.

Current long read sequencing approaches performed transcriptome-wide are inadequate for obtaining sufficient depth of long reads for the targets of interest to accurately quantify the connectivity of AS-APA events. To address this, we developed a probe-based cDNA pulldown strategy to enrich for genes of interest prior to sequencing on the Nanopore platform (Fig. 2a). We call this method Pull-a-Long Seq (PL-Seq). In this approach, SMARTer cDNA synthesis using oligo (dT) priming is performed to obtain full-length cDNA from total RNA, and SMARTer oligo sequence is introduced at 5’ and 3’ ends to enable downstream PCR amplification. Two to five biotinylated probes are designed to enrich each target gene, with the probes targeted to constitutive exons and universal 3’UTR sequences (Supplementary Table 5). After PCR amplification and library preparation, Nanopore sequencing is performed on the nanopore MinION sequencer.

**Figure 2.**
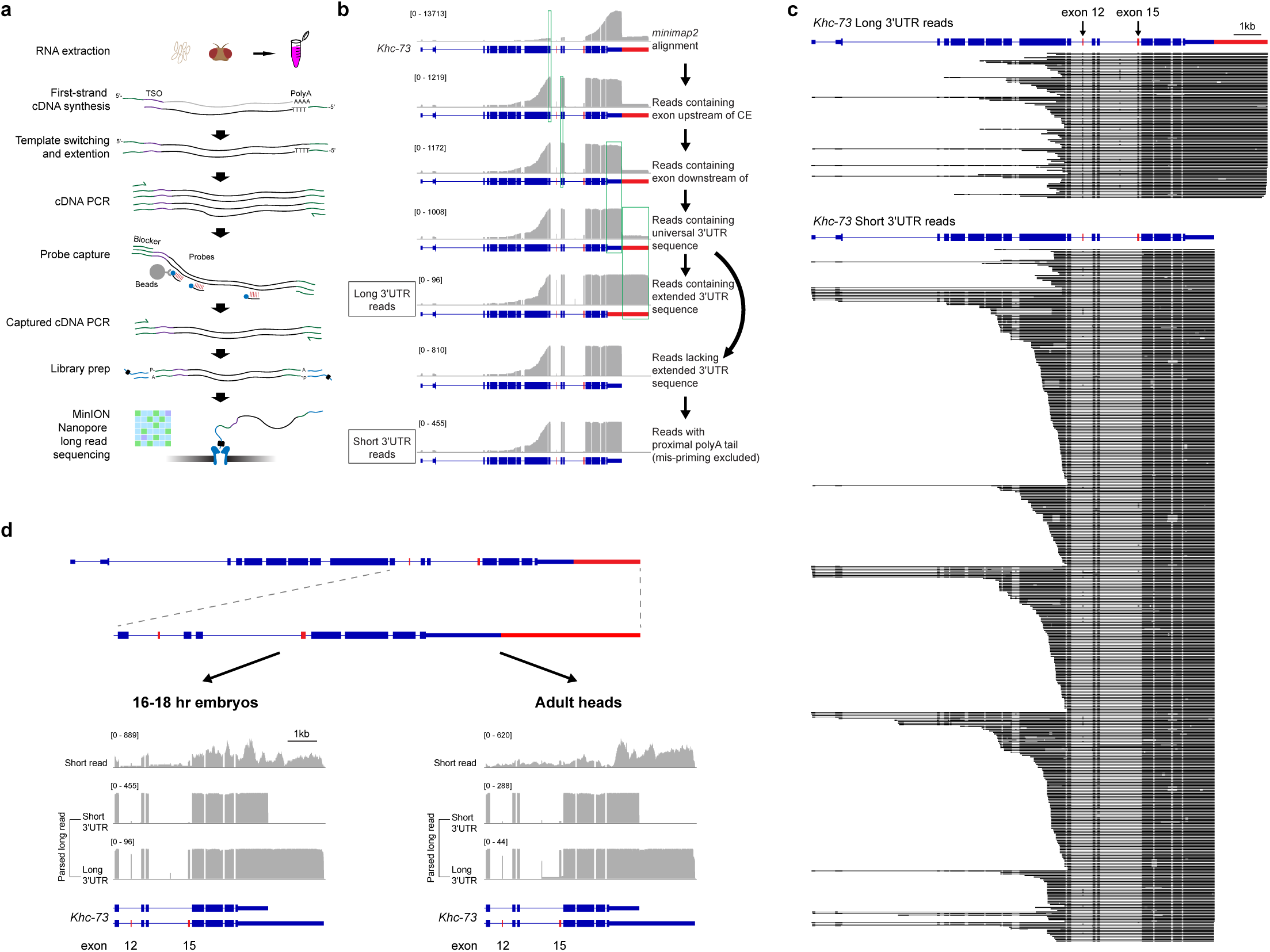
PL-Seq library preparation and data processing workflow. (**a**) Illustration showing PL-Seq library preparation. (**b**) Filtering of reads and parsing to long and short 3’UTR isoforms demonstrated on the *Khc-73* gene. Reads aligned to *Khc-73* are filtered to ensure that they contain upstream and downstream constitutive exons of each specific CE event, and the universal 3’UTR sequence and then parsed to long or short 3’UTR reads. (**c**) Parsed long reads from *Khc-73* long (top) and short (bottom) 3’UTR isoforms in a 16-18 hr embryo sample. (**d**) Coverage tracks from short-read RNA-Seq (top track) and PL-Seq (bottom tracks) showing contrasting CE splicing patterns between parsed short and long 3’UTR reads for *Khc-73* in late-stage embryos (left) and adult heads (right).

A series of alignment and gene-specific filtering steps are required to quantify the upstream exon inclusion for long versus short 3’UTRs. First, alignment to the *Drosophila* genome (dm6) is performed using minimap2^49^, then reads from a given experiment are filtered for regions spanning alternative CEs and alternative 3’ ends for the targeted genes. We outline these filtering steps for the *Khc-73* gene as an example (Fig. 2b). Reads are selected that cover the common 3’UTR region and exons that flank the CE of interest– in this case the reads are required to include the constitutive exons 11 and 13 (Fig. 2b). Reads which span the extended 3’UTR region are selected. These reads serve exclusively as “Long 3’UTR” parsed reads. Then, the remaining reads are filtered to account for mispriming from genomically encoded A stretches, and reads containing untemplated polyA tails are selected to constitute the “Short 3’UTR” parsed reads. Percent-Spliced-In (PSI) values of upstream CEs can then be exclusively assigned to each 3’UTR mRNA isoform. Individual full-length reads from late-stage embryos demonstrates a preferential inclusion of exons 12 and 15 in the long 3’UTR reads compared to the short 3’UTR reads (Fig. 2c, d). This difference is even more striking when observing filtered coverage tracks (Fig. 2d). For both 16-18 hr embryos and adult heads there is a near binary switch in the usage of exons 12 and 15 in long versus short 3’UTR isoforms (Fig. 2c, d). The *Khc-73* long 3’UTR isoform exhibited an exon 15 PSI of 96.8% compared to 1.6% for the short 3’UTR isoform (Fig. 2c). Similar results were obtained from adult heads (Fig. 2d).

To examine the efficiency of the probe-based cDNA capture, we compared embryo PL-Seq libraries prepared with pulldown for 15 genes vs without pulldown. Reads from the 15 targeted genes comprised 0.05% of reads without pulldown. After pulldown, these 15 genes accounted for 46.43% of aligned reads, demonstrating the effectiveness of the enrichment strategy (Fig. 3a). A potential concern with the probe-based pulldown approach is that it could introduce experimental biases that alter PSI value calculation. To test for this possibility, we examined the *stai* gene, which could be sequenced at sufficient depth in the absence of enrichment by cDNA pulldown. *stai* exon 10 PSI was found to not be altered in pulldown versus no pulldown libraries (Fig. 3b). Next, we compared read coverage across gene bodies for the no pulldown versus pulldown library. Both libraries showed a bias for the 3’ end since RT is initiated with oligo dT, but this was more pronounced for the pulldown library (Fig. 3c). Also, a larger portion of novel splicing junctions were observed in pulldown sample versus control library (Supplementary Fig. 4). These were expected given the relatively long length and complexity of the genes targeted for pulldown. Read length distribution was found to be skewed longer in the pulldown library, also reflecting the pulldown of longer cDNAs from the target genes (Fig. 3d).

**Figure 3.**
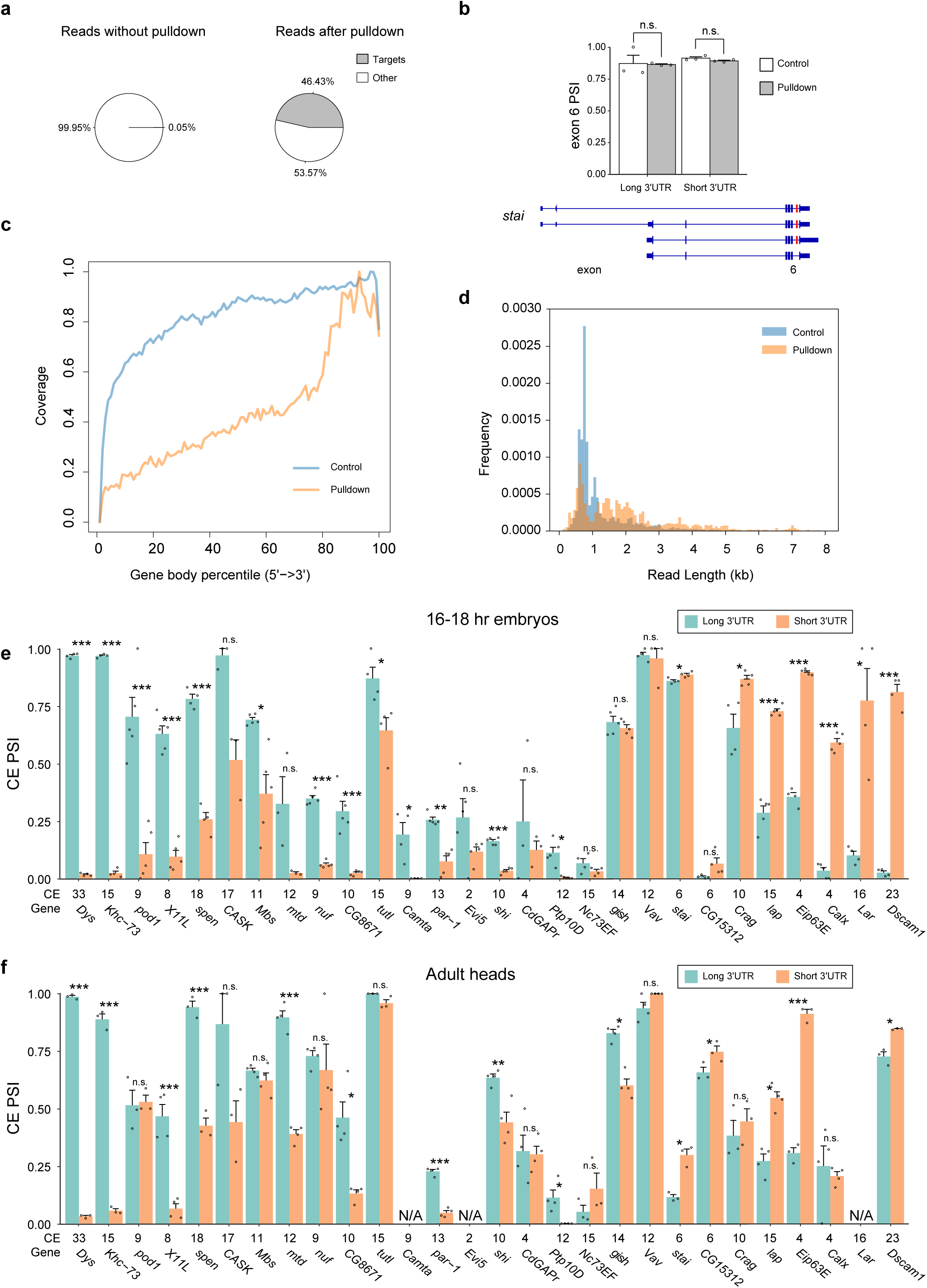
PL-Seq reveals 3’UTR dependent CE splicing for 23 genes. (**a**) Distribution of target gene reads in PL-Seq libraries performed without (left) or with (right) pulldown. (**b**) *stai* exon 10 PSI compared between libraries without pulldown (Control) and with pulldown (Pulldown). For both long and short 3’UTR isoforms, exon 10 PSI shows no significant difference. (**c**) The gene body coverage and (**d**) length distribution of aligned reads in control and pulldown samples. (**e**) CE PSIs for short and long 3’UTR isoforms in late-stage embryos (16-18 hr) from 28 genes. (**f**) CE PSIs for short and long 3’UTR isoforms in adult heads. The cassette exon number for each gene is listed on the x axis (CE). Data is shown as Mean + SEM. * indicates *p*<0.05, ** *p*<0.01 and *** *p*<0.005. n=3-5. Also see supplementary Fig. 4, 5.

We generated two pulldown probe sets that targeted 31 of the 93 genes identified as having regulated AS and APA in late versus early-stage embryos and/or in adult heads versus ovaries (see Supplementary Table 5 for probe density per gene and sequences). PL-Seq data from three to five biological replicates is shown for individual CE PSI values belonging to long or short 3’UTR for 16-18 hr embryos (Fig. 3e) and adult heads (Fig. 3f). Three genes lacked sufficient read depth to quantify changes in 3’UTR specific alternative splicing. Of the remaining 28 genes, we found that 23 genes showed significantly different CE splicing between short and long 3’UTR isoforms either in 16-18 hr embryos (20 genes), adult heads (15 genes), or both (12 genes) (two tailed paired t-test, *p*<0.05). Only 5 of the tested genes with appropriate read coverage showed no significant 3’UTR discrepancy of CE splicing either in 16-18 hr embryos or in adult heads (Fig. 3e, f). For the 23 genes showing connected CE alternative splicing and APA, the CE PSI difference (PSI_long_-PSI_short_) varied widely, from −0.788 to 0.955 (Supplementary Fig. 5). Multiple genes were found to exhibit 3’UTR connected CE splicing even when they were not originally identified as AS-APA genes from short-read data. These included *pod1*, *Crag*, *Eip63E*, *shi*, and *Calx* in embryos, and *Dys* and *Dscam1* in heads (Supplementary Fig. 5). This suggests that 3’UTR connected CE splicing likely affects far more genes than those revealed to have regulated AS and APA in short read RNA-Seq data (Fig. 1e, Supplementary Fig. 1).

PL-Seq revealed interesting cases of connectivity between alternative exons and 3’UTRs. For some genes, trends were different in late-stage embryos compared to adult heads. We previously demonstrated the connectivity of exon skipping events to the long 3’UTR isoform of the *Dscam1* gene using a variety of methods including nanopore sequencing of “long” and “uni” PCR products (as performed for *Khc-73*, see Supplementary Fig. 3a, b)^18^. In late-stage embryos, the *Dscam1* long 3’UTR exhibits 3.7% PSI for exon 23, whereas the short 3’UTR exhibits 79.9% PSI (Fig. 5a). The PSI difference of exon 23 between long and short 3’UTR is reduced in adult heads, but remains significant. In addition, two microexons flanking exon 23 becomes highly expressed and are found to be mainly connected to the long 3’UTR in heads but not embryos. There were several additional genes, including *Khc-73* and *Dys*, that had multiple adjacent CEs included in preferentially in their long 3’UTRs (Fig. 2c, d and Fig. 4b). We previously found that *Dscam1* exon 19 is completely skipped in long 3’UTR isoforms from adult heads, but could not specifically measure this skipping in the short 3’UTR isoform^18^. PL-Seq enabled the detection of short 3’UTR-specific exon 19 PSI for the first time (98.1% in heads) (Supplementary Fig. 6a). As found previously, 0% exon 19 PSI was observed for the long 3’UTR in heads (Supplementary Fig. 6a).

**Figure 4.**
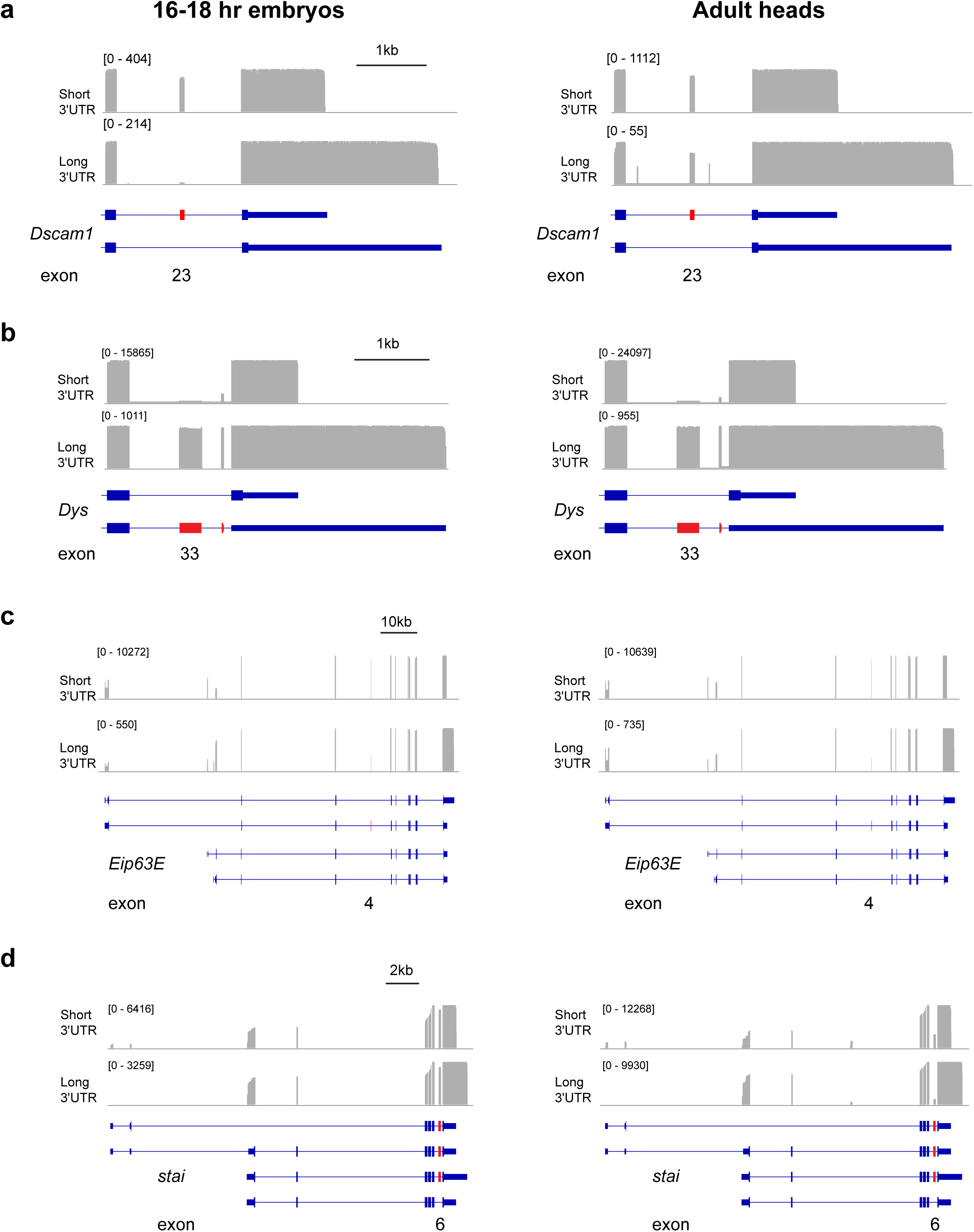
PL-Seq tracks of genes exhibiting 3’UTR connected CE splicing. Left side shows 16-18 hr embryos and right side shows adult heads. (**a**) *Dscam1*, (**b**) *Dys*, (**c**) *Eip63E*, and (**d**) *stai*. For quantification of CE PSI see Fig. 3 and supplementary Fig. 5. Note the change in alternative first exon usage for *Eip63E* and *stai* between long and short 3’UTR isoforms. Also see supplementary Fig. 6.

**Figure 5.**
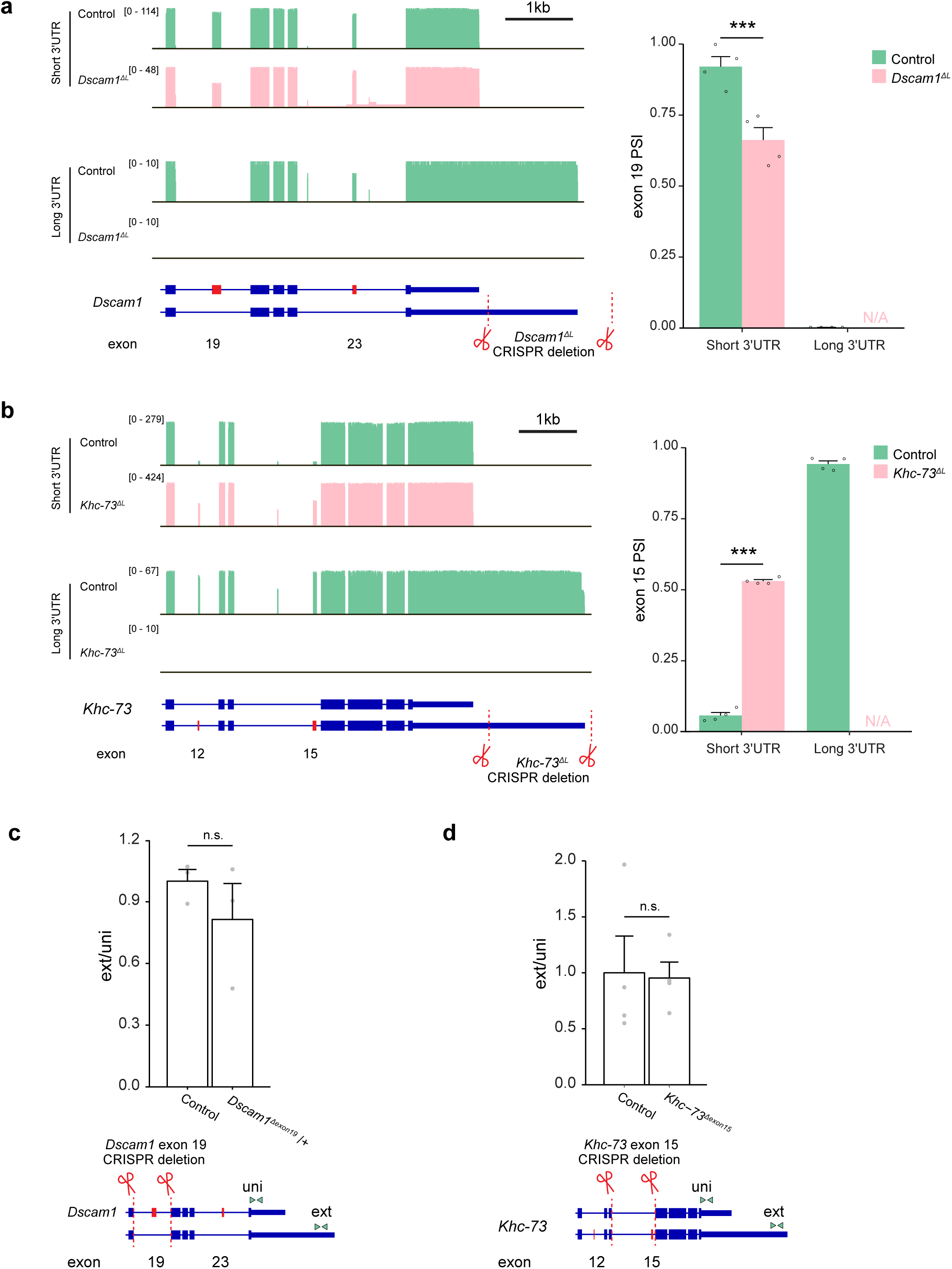
Loss of long 3’UTR affect CE splicing in short 3’UTR isoforms. (**a**) *Dscam1* exon 19 splicing patterns of short and long 3’UTR reads in adult heads from control and *Dscam1* long 3’UTR mutants (*Dscam1^ΔL^*). (**b**) *Khc-73* exon 15 splicing patterns of short and long 3’UTR reads in adult heads from control and long 3’UTR mutants (*Khc-73^ΔL^*). (**c)** qRT-PCR of distal 3’ polyA site usage (ext normalized to uni) in *Dscam1* exon 19 heterozygote mutants (*Dscam1^ΔExon19^*/+). (**d**) qRT-PCR of distal 3’ polyA site usage for *Khc-73* exon 15 mutants (*Khc-73^Δexon15^*). Sequences deleted by CRISPR/Cas9 gene editing are illustrated by dotted red lines and scissors. Approximate location of uni and ext primers are indicated by green arrowheads.

For some genes with relatively shorter full-length sequences (< 4 kb), we were able to obtain an abundance of reads that span the entirety of the gene from 5’ to 3’ end. For example, the *Eip63E* long 3’UTR isoform shows preferential usage of a downstream alternative first exon compared to the short 3’UTR isoform (Fig. 4c). Similarly, the use of an alternative upstream first exon for the *stai* gene occurs for the short 3’UTR isoform but is nearly non-existent in the long 3’UTR isoform (Fig. 4d). PL-Seq analysis revealed other types of 3’UTR connected alternative splicing events beyond CEs. For example, *Calx* exhibits 3’UTR connected mutually exclusive exons (MXE), and these MXEs also behave as alternatively spliced CEs (Supplementary Fig. 6b). We also observed a case for a slight shift in an upstream 3’ splice site depending on 3’UTR choice for *X11L* (Supplementary Fig. 6c), similar to what was found previously for *Dscam1*^18, 50^. Together, these examples demonstrate that PL-Seq reveals the complexity of 3’UTR–linked alternative exon usage.

### Genomic deletion of Long 3’UTR alters CE splicing in short 3’UTR isoforms

In previous work, we found that genomic deletion of the *Dscam1* long 3’UTR (*Dscam1^ΔL^*) altered CE splicing of exon 19 for the remaining mRNAs as measured by RT-PCR^18^. PL-Seq of *Dscam1^ΔL^* fly heads was performed to precisely determine the CE splicing pattern of exons 19 and 23 in short 3’UTR isoforms remaining after long 3’UTR deletion. PL-Seq revealed that *Dscam1^ΔL^* fly heads showed no expression of *Dscam1* long 3’UTR transcripts, as expected (Fig. 5a). In the remaining short 3’UTR mRNA isoforms, there was a significant reduction in exon 19 PSI (Control PSI= 92.1%, *Dscam1^ΔL^* PSI=66.2%, *p*<0.005). The remaining short 3’UTR in *Dscam1^ΔL^* fly heads thus exhibits increased skipping of exon 19. This suggests a feedback system that ensures exon 19 skipped mRNAs are expressed.

We performed a similar CRISPR/Cas9 mediated genomic deletion of the *Khc-73* long 3’UTR (*Khc-73^ΔL^*) to determine whether long 3’UTR loss could impact the exon content of the remaining *Khc-73* short 3’UTR mRNAs. Removal of the genomic region downstream of the proximal polyA site and past the distal polyA site resulted in complete loss of long 3’UTR mRNAs with flies being homozygous viable, as is the case for *Khc-73* null mutants^51^ (Fig. 5b). *Khc-73* short 3’UTR mRNA isoforms normally exhibit very low inclusion of exons 12 and 15, whereas levels of inclusion are much higher in the long 3’UTR mRNAs (Fig. 2d, 5b). In *Khc-73^ΔL^* fly heads the short 3’UTR mRNAs displayed massively increased exon inclusion for exons 12 and 15. Exon 15 PSI in the short 3’UTR mRNA was increased almost 10-fold from 5.4% to 52.7% (*p*<0.001). These PSI values approached what was found for exon 15 PSI in the WT long 3’UTR samples. Thus, both *Dscam1* and *Khc-73* short 3’UTRs exhibit alteration in CE AS upon genomic loss of the long 3’UTR region. These results suggest that alternative splicing of these exons might be influenced by 3’ end processing or the sequence content of long 3’UTRs.

Most splicing events are considered to be spliced co-transcriptionally^52^; thus, an alternative hypothesis is that the strong connectivity of CEs with alternative 3’UTRs is dictated by the CE event regulating downstream APA. To test this, we forced the removal of CEs in *Dscam1* exon 19 and *Khc-73* exon 15 by deleting these exons and their flanking introns. *Dscam1* exon 19 deletion was found to be homozygous lethal; thus, we performed analysis on heterozygous mutant flies. The *Khc-73* exon 15 deletion flies were homozygous viable. We measured the impact of these deletions on the relative expression of the long to short 3’UTR by RT-qPCR and for both we found this to be unchanged (Fig. 4c, d). This suggests that CE alternative splicing does not impact 3’UTR choice for *Dscam1* and *Khc-73*.

### 3’UTR connected CE alternative splicing is deregulated in *elav*^5^ mutants

ELAV and the related protein FNE are key regulators of AS and APA in the nervous system^10, 13, 18^. We re-analyzed short-read RNA-Seq data from L1 CNS samples of *elav,fne* double mutants versus control flies to obtain AS and APA changes^9^ (Supplementary Fig. 7a, b). We identified 29 genes that are regulated by CE alternative splicing and 3’UTR shortening in the mutant condition compared to wild type. The 3’UTR shortening in these mutants was significantly associated with upstream CE alternative splicing regulation (Fisher’s exact test, *p*=0.0004, Fig. 6a). Out of the 29 ELAV/FNE regulated AS-APA genes, 3’UTR connected PSI data was available for 10. Eight out of 10 genes showed 3’UTR dependent CE alternative splicing in late-stage embryos and/or adult heads (Fig. 6b). 23 genes have been shown to have both regulated AS and APA in earlier stage embryos of *elav,fne* mutants using different analysis methods^13^. Our PL-Seq data collected from late-stage embryos and adult heads included seven of them while six show 3’UTR dependent CE splicing (Supplementary Table 8). We reasoned that PL-Seq could be used to determine if CE splicing regulated by ELAV/FNE somehow depend on which 3’UTR isoform is selected downstream. *elav,fne* double mutants would have long 3’UTR isoform expression near zero for many genes, making it extremely difficult to quantify CE alternative splicing in long 3’UTR isoforms. In contrast, from L1 CNS samples of *elav*^5^ mutants display much fewer APA changes^9^ (Supplementary Fig. 7c, d). Thus, we performed experiments with *elav*^5^ mutant embryos instead, as they retain some expression of long 3’UTRs.

**Figure 6.**
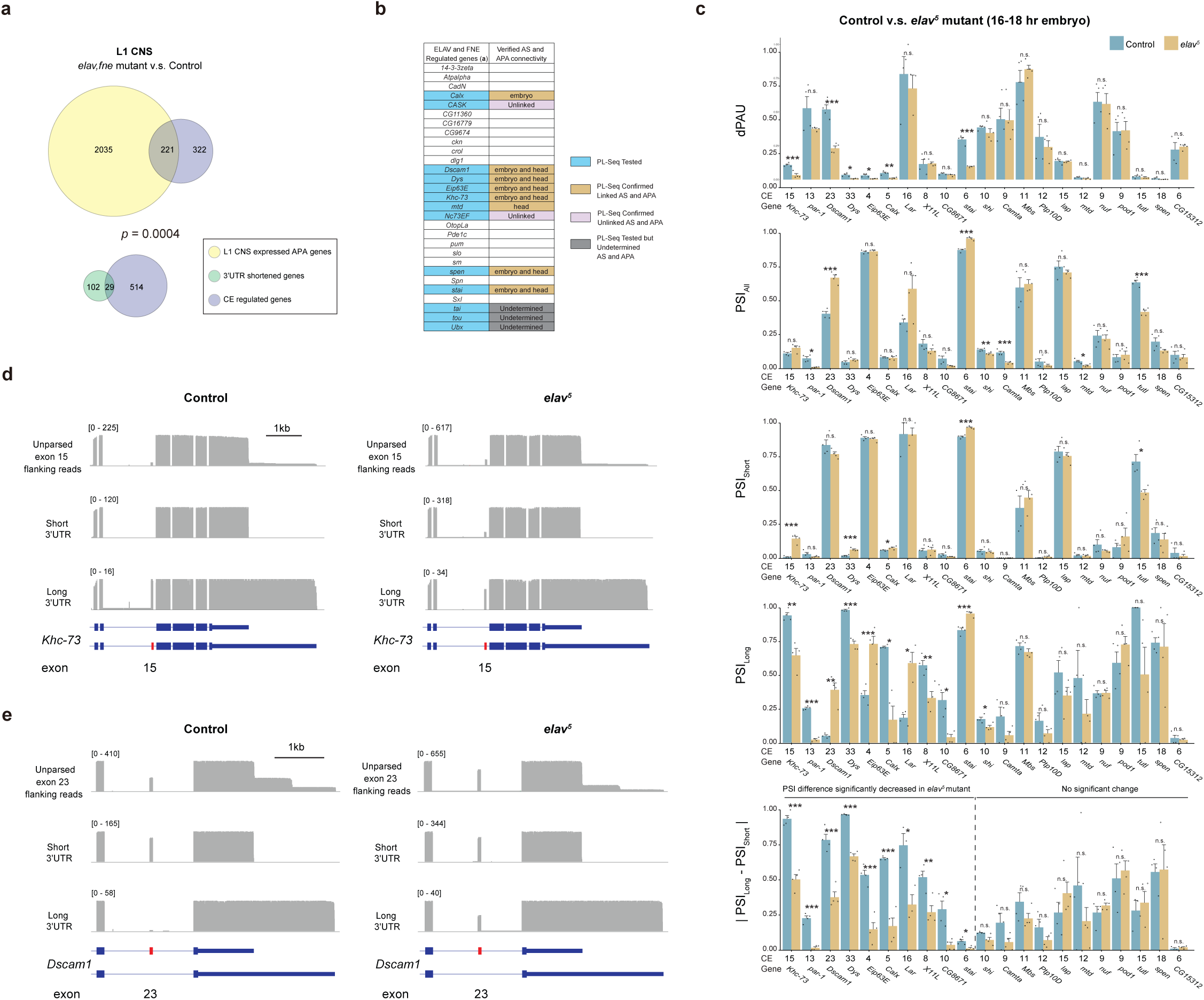
ELAV loss of function deregulates 3’UTR connected CE splicing. (**a**) Fisher’s exact test shows 3’UTR shortening genes in *elav,fne* mutants are significantly associated with regulated CE events when comparing L1 CNS *elav,fne* mutants with controls. Data re-analyzed from published work^1^. (**b**) List of the 29 AS and APA regulated genes in *elav,fne* mutants with PL-Seq verification information for 3’UTR connected CE splicing. (**c**) dPAU, PSI from all transcripts (PSI_All_), short (PSI_Short_) and long (PSI_Long_) 3’UTR isoforms, and CE splicing difference between short and long 3’UTR isoforms (|PSI_Long_ - PSI_Short_|) compared between control and *elav* mutant embryos using PL-Seq data. Data is shown as Mean + SEM. * indicates *p* < 0.05, ** *p*<0.01 and *** *p*<0.005. Two tailed t-test, n=3-4. (**d**) *Khc-73* exon 15 splicing patterns of short and long 3’UTR reads in adult heads from control and *elav* mutant embryos. (**e**) *Dscam1* exon 23 splicing patterns of short and long 3’UTR reads in adult heads from control and *elav* mutant embryos. Also see Supplementary Fig. 7.

To investigate the role of ELAV in APA and CE AS connectivity, we performed PL-Seq for the previously validated 23 AS-APA genes and several control genes to monitor the *elav* mutation (Supplementary Table 5). PL-Seq data collected from *elav* mutant and control embryonic samples revealed significant changes of: (1) distal polyA site usage (dPAU) for 6 genes, (2) CE splicing irrespective of 3’UTR isoform (PSI_All_) for 7 genes, (3) CE splicing in short 3’UTR isoform only (PSI_Short_) for 5 genes, (4) CE splicing in long 3’UTR isoforms (PSI_Long_) for 11 genes (Fig. 6c).

For *Khc-73*, we observed a significant reduction in distal polyA site usage (dPAU) in the *elav* mutant condition (Fig. 6c, e). In *elav* compared to control embryos, exon 15 PSI decreased for the long 3’UTR isoform and increased for the short 3’UTR isoform. In contrast, when measuring exon 15 PSI from all aligned reads, there was no change in *elav* mutants compared to control (Fig. 6c, e). Thus, the direction of ELAV-mediated PSI change was dependent on 3’UTR isoform. Interestingly, PL-Seq revealed that the increased retention of *Dscam1* exon 23 in *elav* mutant embryos occurred exclusively in the long 3’UTR whereas there was no change in the short 3’UTR transcripts (Fig. 6d). This evidence from *Khc-73* and *Dscam1* suggest that 3’UTR isoform content or choice impacts ELAV-mediated alternative splicing of CEs. For both *Dscam1* and *Khc-73*, there was a significant reduction in the difference of PSI in long versus short 3’UTR isoforms in *elav*^5^ embryos compared to control (control vs *elav*^5^ |PSI_Long_-PSI_Short_|, *p*<0.001) (Fig. 6c). Remarkably, for all significant changes in |PSI_Long_-PSI_Short_| detected (10/23 genes), there was always a decrease in the *elav*^5^ condition (Fig. 6c). In other words, after controlling for the directionality of inclusion/skipping, the loss of ELAV tends to minimize the difference in 3’UTR mRNA isoform specific CE PSI values. Overall, these data illustrate the utility of PL-Seq to quantify RNA binding protein-regulated changes in the intramolecular connectivity of 3’UTRs to CEs.

## Discussion

AS and APA are key co/post-transcriptional processing events impacting most metazoan genes. Despite their importance, a limited number of studies have found evidence that alternative exon choice and 3’UTR choice are connected.^18, 28^ Here, we used PL-Seq, a cDNA capture based long-read sequencing method, to investigate the interactions between AS and APA. In *Drosophila* late-stage embryos and heads, we uncovered 3’UTR connected CE splicing events for 23 genes, of which 10 were not initially recognized as potential candidates using short-read RNA-Seq analysis. These findings suggest that many more genes might be affected by connected AS-APA events.

To obtain a broader scope of 3’UTR linkage to CE splicing, PL-Seq could be applied to all 694 genes expressed in *Drosophila* embryos/heads that are annotated for both CEs and alternative length 3’UTRs. Outside of the nervous system, there are surely other connected AS-APA events waiting to be discovered in different tissues, developmental time points, disease states, and cell types exhibiting regulation of APA^1, 53^. We restricted our sequencing to explore CE alternative splicing, but there is likely a plethora of exon to 3’ end connectivity that await discovery with long read sequencing. For example, alternative promoter usage has been previously connected to alternative polyA site usage^54^, and enhancer activity/transcriptional activity has been shown to regulate APA^55, 56^. We identify several examples of such coordination here (Fig. 4c, d). However, investigating the scope of these events could be challenging due to the often large distance between first and last exons.

cDNA capture-based nanopore sequencing methods such as PL-Seq are valuable additions to the transcriptomics tool-box^43, 44^. Many of the limitations of read length and depth associated with long-read technologies applied transcriptome-wide should eventually be overcome as these sequencing approaches mature. Until then, PL-Seq is a particularly useful approach for quantifying the alternative exon content of alternative 3’UTR mRNA isoforms. PL-Seq is low-cost and requires little to no capital investment. The method and analysis pipeline can be applied to quantify exon to 3’UTR connectivity for specific genes of interest by any laboratory equipped for molecular biology research. While we used a limited number of probes and targets in our current work, capture-based cDNA pulldown is effective at enriching thousands of targets simultaneously and can be scaled up accordingly^41, 42, 44^. PL-Seq should also be adaptable to single-cell analysis, providing a targeted approach to complement recent advances in long-read single cell RNA-seq^28, 57^.

Tandem 3’UTR APA events by definition do not alter the protein-coding potential of the mRNA isoforms produced, in contrast to intronic APA or alternative last exon APA. However, for certain genes with extreme connectivity of upstream CEs to tandem 3’UTRs, tandem 3’UTR APA choices are inseparable from the production of different protein isoforms (e.g. *Dys*, *Khc-73, Dscam1*). Such examples do not fit neatly into our current classifications systems of alternative splicing and alternative polyadenylation. In most of the 23 AS and APA connected genes we study here, the differences between the isoforms are only in the amino acids encoded in the CEs. However, in other cases, CE inclusion can cause a frameshift that changes the C-terminus of the protein more drastically. For instance, the inclusion of CEs in the *Dys* long 3’UTR isoform leads to the stop codon shifting to the second last exon and eliminates the protein coding capacity of the terminal exon (Fig. 4b).

To explore the mechanism of intramolecularly connected AS and APA, we generated CRISPR deletion mutants lacking either long 3’UTR isoforms or inclusion isoforms of specific CEs. Previous studies have shown that splicing can affect 3’ end processing^36, 58, 59^, but our results did not provide any evidence that loss of alternative splicing of an upstream CE affects alternative polyA site choice for *Dscam1 and Khc-73* (Fig. 5c,d). However, we did observe that the loss of long 3’UTR isoforms resulted in a change in the CE splicing pattern in short 3’UTR isoforms, which we quantified using PL-Seq (Fig. 5a, b). The mechanism of how loss of long 3’UTR impacts CE splicing for genes like *Dscam1* and *Khc-73* is still unclear. Future investigation into the timing of 3’UTR transcription and processing relative to splicing, using nascent RNA technologies coupled to long read sequencing^36, 39^, might lend new insights into how ELAV and 3’UTR sequence can impact upstream AS events.

## Acknowledgements

We thank Dr. Christopher Vollmers (UC Santa Cruz) for Nanopore sequencing methodology discussions and Dr. Jung Hwan Kim for insights and discussion on the manuscript. Thanks to Miura lab members for reading and providing input on the manuscript. This work was supported by NSF IOS grant 1656463 and NIGMS grant R35 GM138319 awarded to Dr. Miura. Core facilities at the University of Nevada, Reno campus were supported by NIGMS COBRE P30 GM103650.

## Author Contributions

Conceptualization, Z.Z., W.C., B.B., and P.M; Methodology, Z.Z., W.C., B.B., and P.M.; Investigation, Z.Z., B.B, and W.C.; Data Analysis, Z.Z., B.B, W.C., and P.M. Software, Z.Z.; Writing, Z.Z., and P.M.; Funding Acquisition, P.M.; Supervision, P.M.

## Methods

### *Drosophila* CRISPR/Cas9 deletion lines

Fly CRISPR/Cas9 genome editing was performed by WellGenetics Inc. A homology-directed repair strategy was utilized to generate flies harboring a deletion of the *Khc-73* long 3’UTR sequence Briefly, two gRNAs were designed targeting the long 3’UTR and sequence from chr2R:15515657 to chr2R:15517432 was deleted. The donor plasmid containing the homologous arms with the deletion and two loxP sites bracketed 3xP3-RFP cassette was injected into embryos of control strains together with targeting gRNAs. Flies carrying RFP marker were selected out and further validated by genomic PCR and sequencing. Validated mutants were crossed to flies expressing Cre recombinase constitutively to remove 3xP3-RFP insertion.

For CE mutants, CE including the flanking introns was deleted using the homology-directed repair strategy. 3xP3-DsRed flanked by PiggyBac terminal repeats was embedded in a TIAA motif in the homologous sequences to avoid additional restriction digestion residues. For *Khc-73* exon 15 mutant sequence from chr2R:15520337 to chr2R: 15521841 was deleted, and for *Dscam1* exon 19 mutant sequence from chr2R:7323333 to chr2R: 7324569 was deleted. DsRed was used as the screening marker and then excised from validated mutant flies by crossing to flies expressing the PiggyBac transcriptase.

### Short-read RNA-Seq data based alternative splicing and alternative polyadenylation analysis

For AS and APA analysis, we used publicly available RNA-Seq data from fly tissues under BioProject accession PRJNA75285. For AS analysis, RNA-Seq reads were aligned to the *Drosophila Melanogaster* (dm6) genome using STAR 2.7. Sorted output bam files were then fed into rMATS (v4.0.2)^48^ to identify alternatively spliced CEs. The output file was filtered by FDR < 0.05 and | IncLevelDiff | > 0.2 to create a gene list of differential spliced events of high confidence. For alternative polyadenylation analysis, QAPA 1.2.3 was used with Ensembl gene dm6 3’UTR annotations. Custom R scripts were used for filtering as follows: only genes with gene-level TPM values greater than 1 were considered as expressed and thus included in the downstream analysis; dPAU values were set by filtering the maximum length of the 3’UTR sequence detected by QAPA; fold change of dPAU values between samples was calculated and t-test was performed followed by FDR correction; genes that were differentially regulated at both AS and APA levels were identified (fold change of dPAU>2 or <0.5 plus FDR<0.05), and their association was tested by Fisher’s exact test. Gene Ontology analysis was performed on FlyEnrichr^60^. Data arrangement, statistical analysis, and graph generation was performed in R 3.6.1 and R 4.2.2.

### RNA extraction, cDNA synthesis, nested RT-PCR and qRT-PCR

Fly embryos from various time points and adult heads from mixed 1-4-day old males and females were collected. Total RNA was extracted using Trizol (Thermo Fisher Scientific) per the manufacturer’s instructions. Briefly, samples were triturated and lysed in Trizol for 10 min at room temperature. Phase separation was performed by adding 1/5 volume of chloroform and centrifuging at 20,000xg for 20 min at 4°C. Upper aqueous phase was precipitated with isopropanol and centrifuged at 20,000xg for 20 min at 4°C. RNA pellet was washed with 70% ethanol (ethanol was removed after centrifugation at 20,000xg for 10 min at 4°C), and then resuspended in desired volume of distilled water. RNA was quantified using a Nanodrop spectrometer.

For cDNA synthesis, 1 ug of Turbo DNase (Thermo Fisher Scientific) treated total RNA was reverse transcribed using Maxima reverse transcriptase (Thermo Fisher Scientific). For nested RT-PCR, the first round of PCR was performed using LongAmp Taq 2X master mix (NEB) and 2 – 3 uL of cDNA (1:5 diluted in water) for 30 cycles per manufacturer’s instructions. PCR products were gel purified using QIAquick gel extraction kit (Qiagen) and eluted in 50 uL of water. Eluted DNA was quantified using a Nanodrop spectrometer. For the second PCR, Taq DNA polymerase with standard buffer (NEB) was used with 25 cycles. PCR products were resolved in 2.5-3% agarose gel and imaged using Gel Doc EZ (Bio-Rad). Exposure time was adjusted to ensure band intensities were not saturated. PSI values were estimated using a gel analyzer tool in Image Lab Software (Bio-Rad). For qRT-PCR, 2 uL of 1:10 diluted cDNA (in water) was used as the template, 1 uL of forward and reverse primers, 10 uL of SYBR Select Master Mix (Thermo Fisher Scientific) and 7 uL water were added to each reaction. Samples were subjected to 40 amplification cycles, and data was collected and analyzed using CFX Maestro Software (Bio-Rad).

### PCR-based Gene-specific Nanopore sequencing

For PCR based Nanopore library preparation, PCR amplicons using *Khc-73* specific primers with barcoding adapter sequences were used. Each sample was barcoded by PCR using Nanopore PCR barcoding kit (EXP-PBC001). Barcoded samples were pooled at equimolar concentration, and end-prepped using NEBNext FFPE DNA Repair Mix and NEBNext Ultra II End Repair Kit. The nanopore adapter was ligated using Nanopore ligation sequencing kit (SQK-LSK109). Alternatively, samples were prepared without barcoding and sequenced separately. In this case, PCR amplicons at the equimolar concentration were end-prepped and the nanopore adapter was directly ligated. MinION Mk1B device and FLO-FLG001 flow cells were used for sequencing of the libraries.

### PL-Seq library preparation

For full-length cDNA synthesis, SMARTer PCR cDNA synthesis kit (Clontech) was used according to the manufacturer’s specifications. In short, total RNA was Dnase treated on-column using PureLink Dnase (Thermo Fisher Scientific) and the PureLink RNA Mini kit (Thermo Fisher Scientific). First strand synthesis was performed using ∼500ng of Dnase treated RNA, 3’ SMART CDS Primer II A, and SMARTer II A TSO. cDNA was diluted 1:5 in TE buffer, and then used as the template to synthesize double strand cDNA amplicons by PCR (optimal 17 – 21 cycles) using Advantage 2 PCR kit according to the manufacturer’s specifications (Clontech). cDNA was purified using NucleoSpin Gel and PCR Clean-up kit (Takara Bio). cDNAs of our interest were enriched by pulldown starting with 5-10 ug PCR synthesized cDNA and custom designed 5’ biotinylated oligonucleotide xGen Lockdown probes (Integrated DNA Technologies) and the xGen hybridization and wash kit (Integrated DNA Technologies). Probe sequences are listed in Supplementary Table 5. Captured cDNA was amplified using Takara LA Taq DNA polymerase Hot-Start version (Clontech) and purified using 1:1 (vol: vol) AMPure XL beads (Beckman Coulter). cDNA prepared as above was end-prepped using NEBNext Companion Module for Oxford Nanopore Technologies Ligation Sequencing (NEB) and the nanopore adapter was ligated using Nanopore ligation sequencing kit (SQK-LSK110). Thirty uL of the prepared library was then loaded on to a flow cell (FLO-FLG001) and sequenced using Nanopore MinION Mk1B sequencer.

### PL-Seq data read mapping, filtering, and analysis

Reads were firstly aligned to fly genome assembly (dm6) and transcriptome using minimap2 (v2.17) with the arguments -ax splice -B 3 -O 3,20 to allow optimized splicing junction recognition. Aligned files from minimap2 were converted to bam format using samtools (v1.6)^61^ and then quality check was performed using tools in NanoPack (v1.41.0)^62^ and coverage was examined using RSeQC (v5.0.1)^63, 64^. Reads aligned to targeted genes were counted by featureCounts^65^. For downstream analysis, aligned read were subjected a series of sequential filtering using a custom python script exon_coverage.py (https://github.com/markandtwin/Pull-a-long). This allows for only including reads that cover upstream constitute exons and downstream universal 3’UTR regions with subsequent parsing for short and long 3’UTR isoforms. The 3’ ends were inferred by drops in Nanopore read coverage in the 3’UTR region and current Ensembl genes annotations. Reads identified as short 3’UTR isoform specific were fed to another custom python script polyA_filtering.py to filter out truncated reads lacking the polyA tail in the soft clipped region and reads that are internally misprimed due to genomically encoded A enriched sequences in the transcripts. Minimum reads count required for PSI calculation from each isoform is 2. PSI values of short and long 3’UTR isoforms were calculated by a third script calculate_PSI.py and tested by pairwise t-test with 3 or more replicates. When three or more tandem polyA sites were used in the gene, long was defined as the most distal 3’UTR and short as the most proximal one. Details for the 3’UTR connected CE we used for the analysis can be found in Supplementary Table 2.

### Data Availability

All the sequencing data is deposited as SRA Bioproject PRJNA771049.

### Code Availability

The custom scripts for PL-Seq workflow are available at: https://github.com/markandtwin/Pull-a-long.

## Supplementary Figures

**Supplementary Figure 1.**
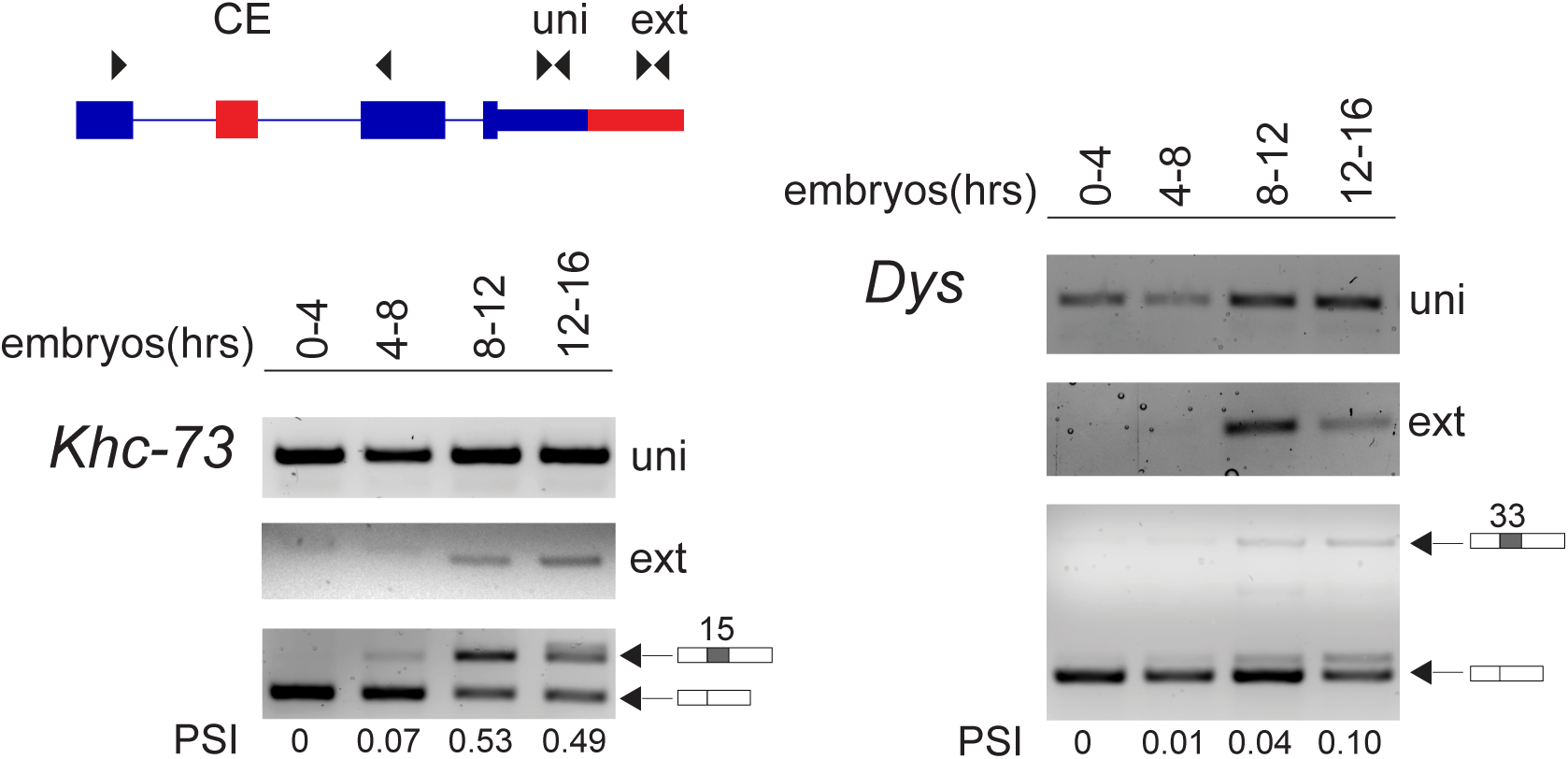
RT-PCR confirms trend of coordinated CE splicing and 3’UTR lengthening during embryonic development.

**Supplementary Figure 2.**
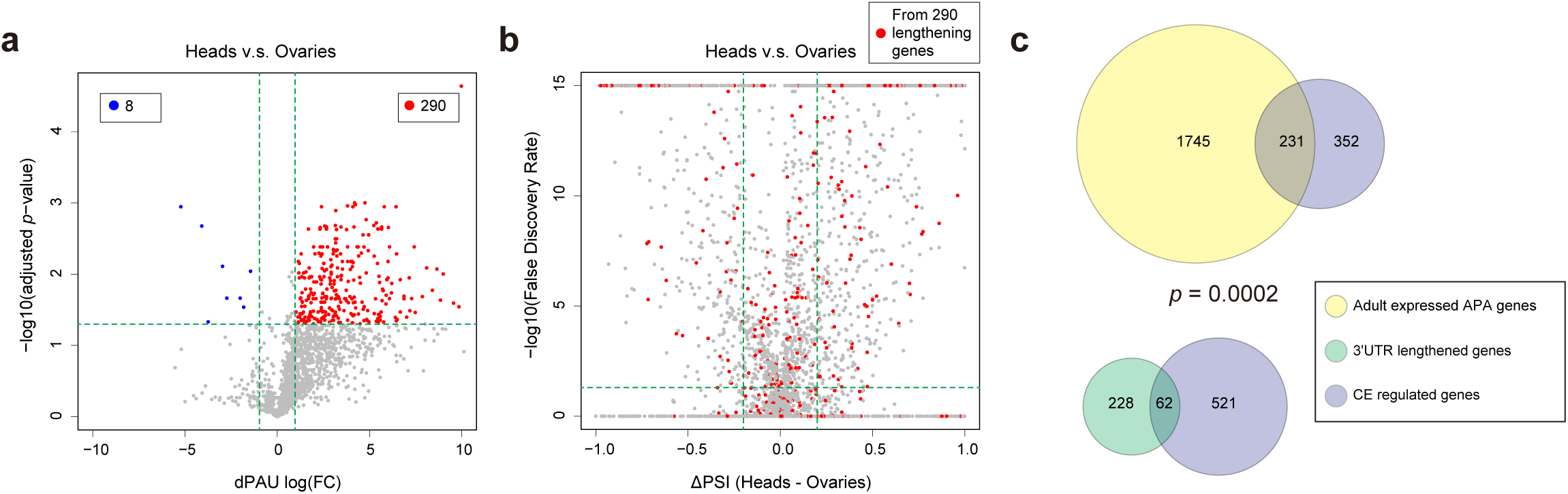
CE AS is associated with 3’UTR lengthening in adult heads. (**a**) Short read RNA-Seq analysis of adult head versus ovary shows that 290 genes exhibit significant 3’UTR lengthening. (**b**) Distribution of PSI change (PSI (heads) – PSI (ovaries)) of adult head vs ovary samples. From the 290 3’UTR lengthening genes (red dots), 63 significant CE skipping and 127 inclusion events are revealed. (**c**) Fisher’s exact test shows that in adult head vs ovary samples, 3’UTR lengthening genes are significantly associated with regulated CE events. Horizontal dash lines in (a) indicate adjusted *p*=0.05, and in (b) indicate adjusted FDR=0.05. Vertical dash lines in (**a**) indicate FC=0.5 (left) and FC=2 (right), while in (b) indicate ΔPSI=-0.2 (left) and ΔPSI =0.2 (right).

**Supplementary Figure 3.**
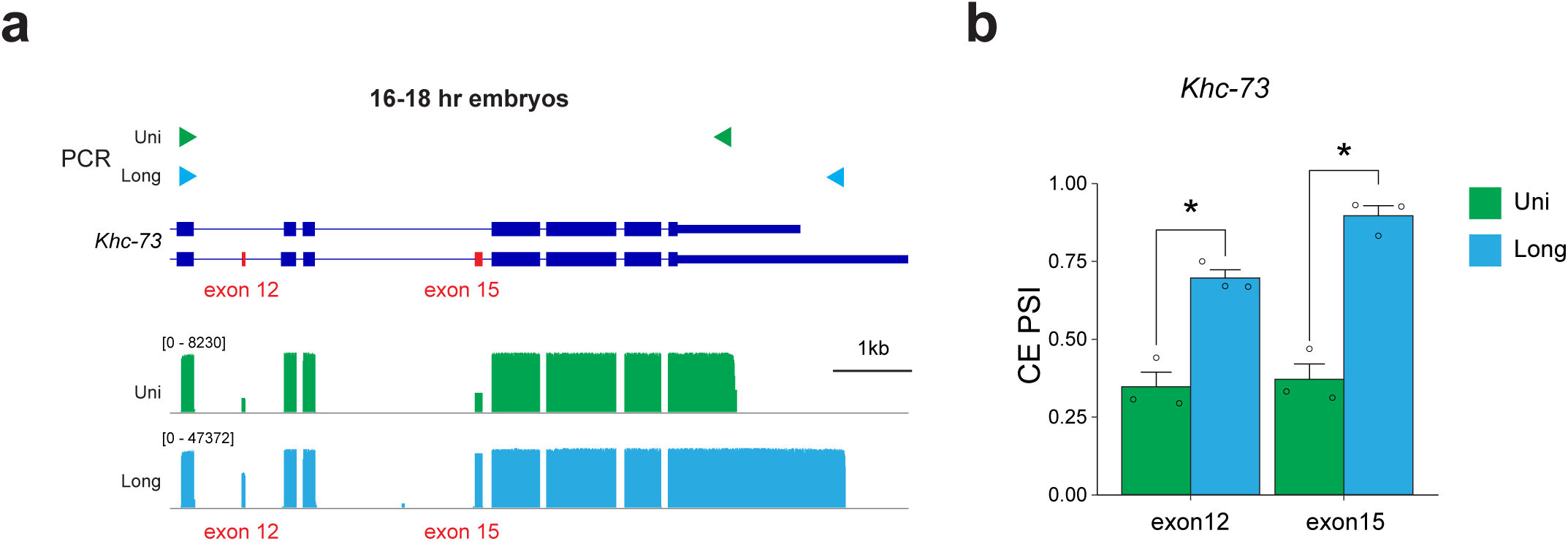
Resolving CE patterns between 3’UTR isoforms using targeted PCR based Nanopore sequencing. (**a**) RT-PCR amplification strategy for performing Nanopore sequencing of targeted region of *Khc-73* (top). Coverage plot of long-reads are shown corresponding to the amplicons that cover all isoforms (Uni) or long 3’UTR isoform specifically (Long). (**b**) Bar plot representing PSI calculated from Nanopore long-read sequencing of long and uni samples. Mean+SEM is shown, n=3 biological replicates. Paired two-tailed t-test. * indicates *p*<0.05.

**Supplementary Figure 4.**
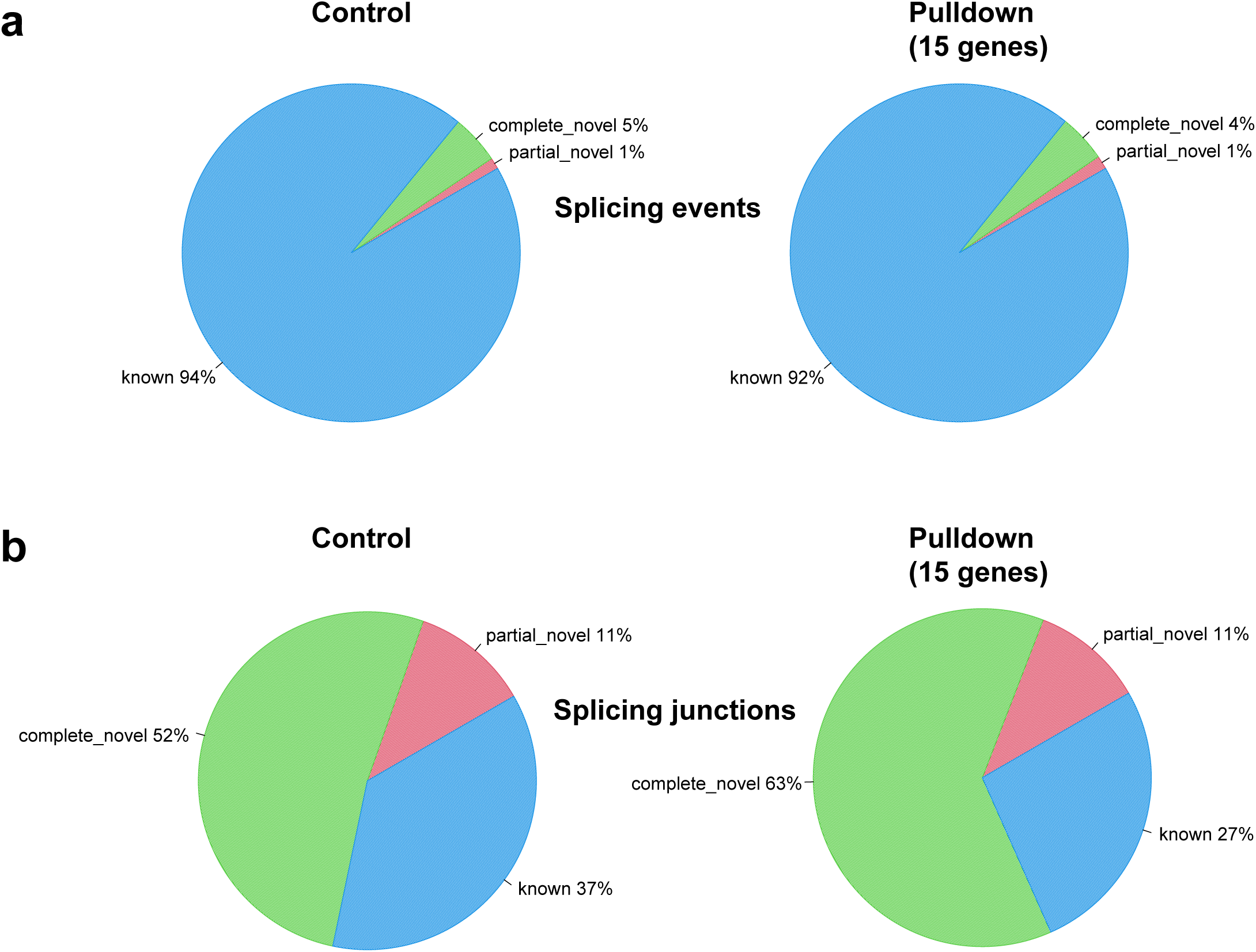
Distribution of splicing events detected from PL-Seq. Pie charts displaying read distribution for (a) Splicing events and (b) Splicing junctions for libraries without pulldown (Control) and with pulldown (Pulldown). Reads are characterized as complete novel, partial novel or known according to the fly Ensemble dm6 annotation.

**Supplementary Figure 5.**
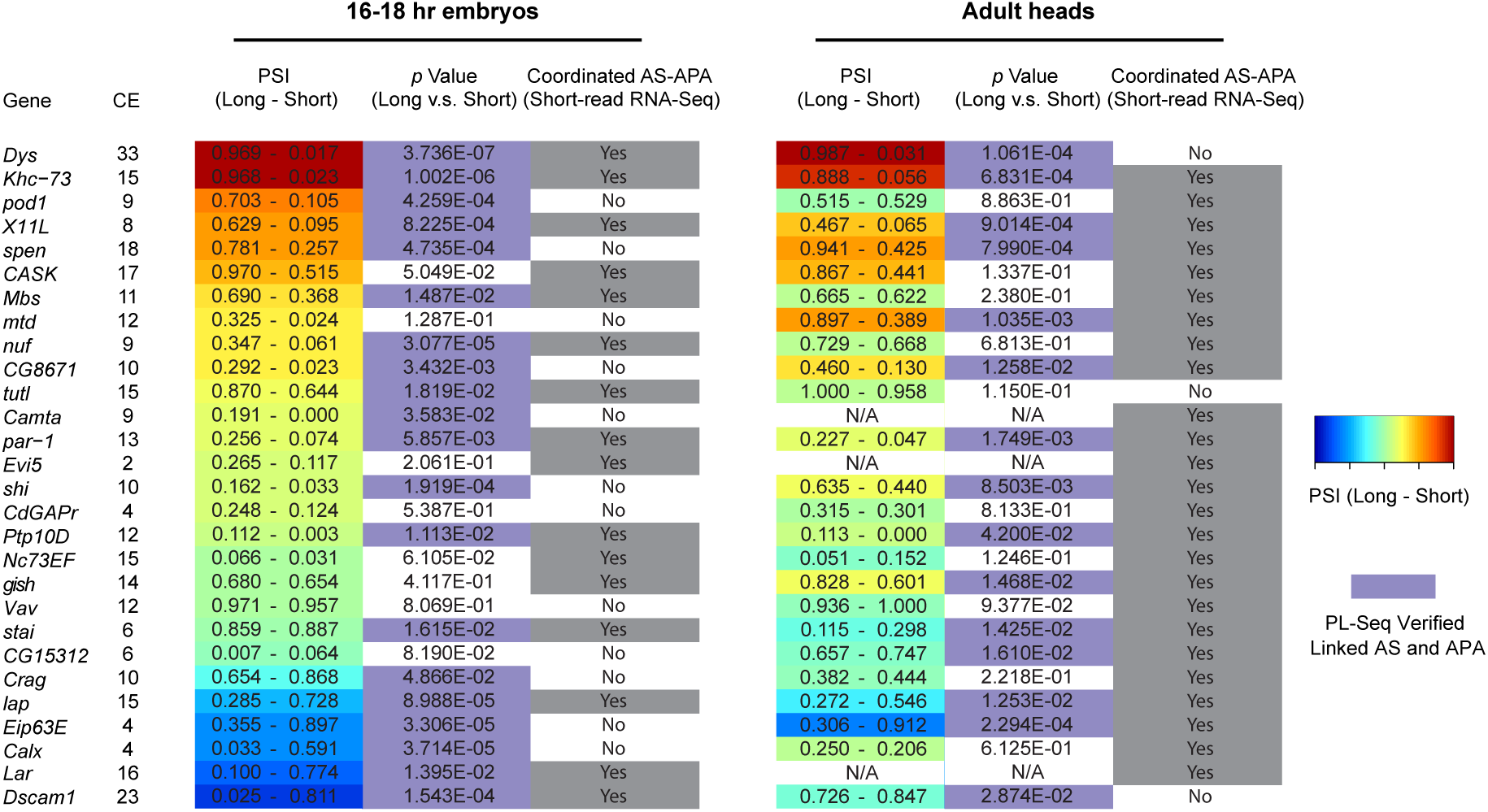
Summary of PSI differences between long 3’UTR and short 3’UTR isoforms. CE splicing difference derived from PL-Seq analysis is shown as PSI (long – short) by heatmap. Individual values of PSI from Long 3’UTR and Short 3’UTR isoforms are listed in the colored cells. *p* values from two tailed paired t-test using PL-Seq data are shown. Purple coloring indicates a significant difference in the t-test. “Coordinated AS-APA” (“Yes” or “No”) refers to whether the gene was determined to exhibit regulated AS and APA from short read RNA-Seq analysis (Fig. 1d for Late vs Early stage embryos, and Supplementary Fig. 1b for Heads vs Ovaries).

**Supplementary Figure 6.**
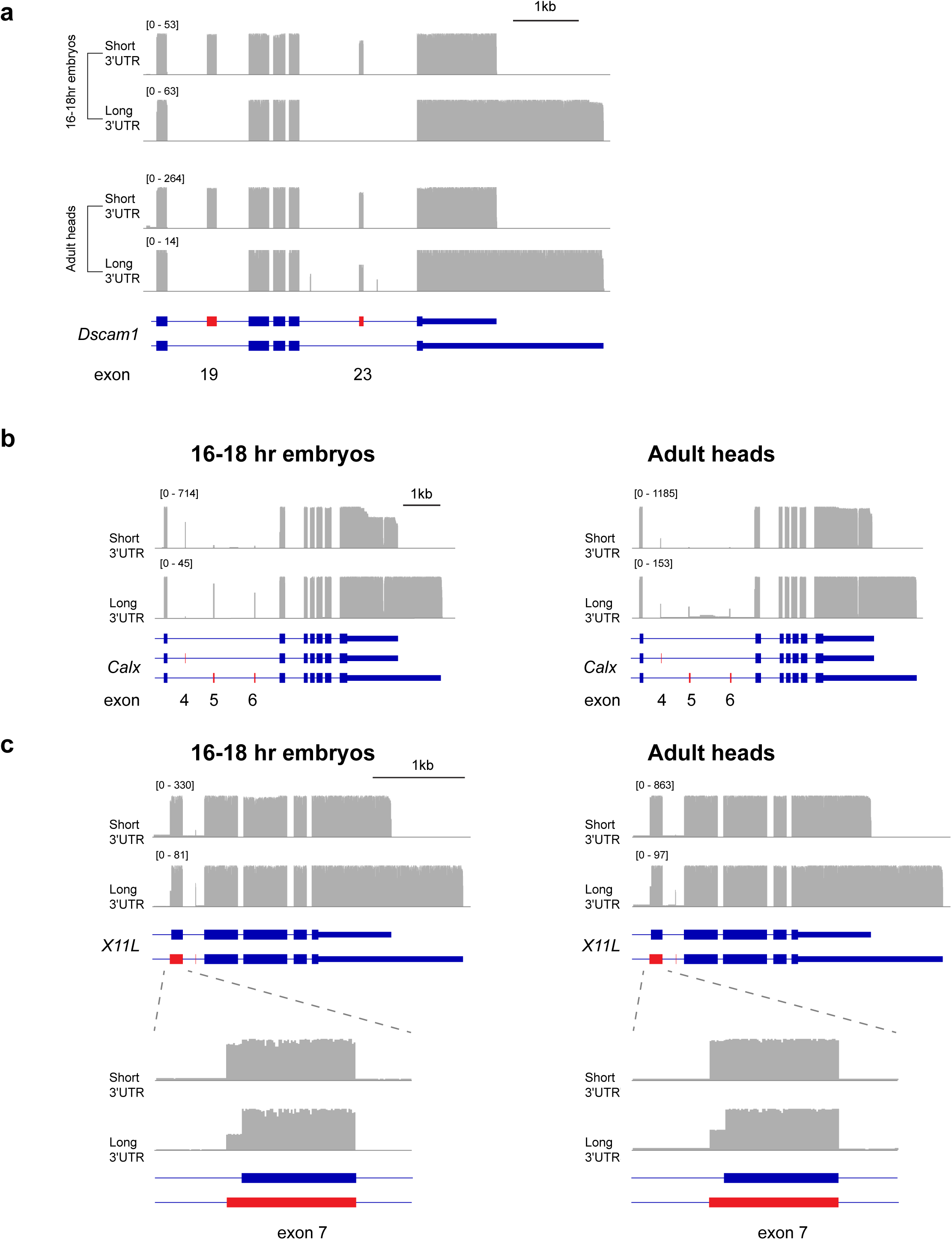
Additional PL-Seq data tracks of genes exhibiting 3’UTR connected AS. (**a**) Re-filtering of reads to detect *Dscam1* exon 19 AS in 16-18 hr embryos and adult heads. (**b**) Coverage tracks of PL-Seq data showing splicing pattern in short and long 3’UTR reads. For *Calx*, exon 3 acts as both a CE and mutually exclusive to exon 4 and 5. (**c**) For *X11L*, exon 8 AS and the usage of an alternative upstream 3’ splice site of exon 7 in short vs long 3’UTR isoforms.

**Supplementary Figure 7.**
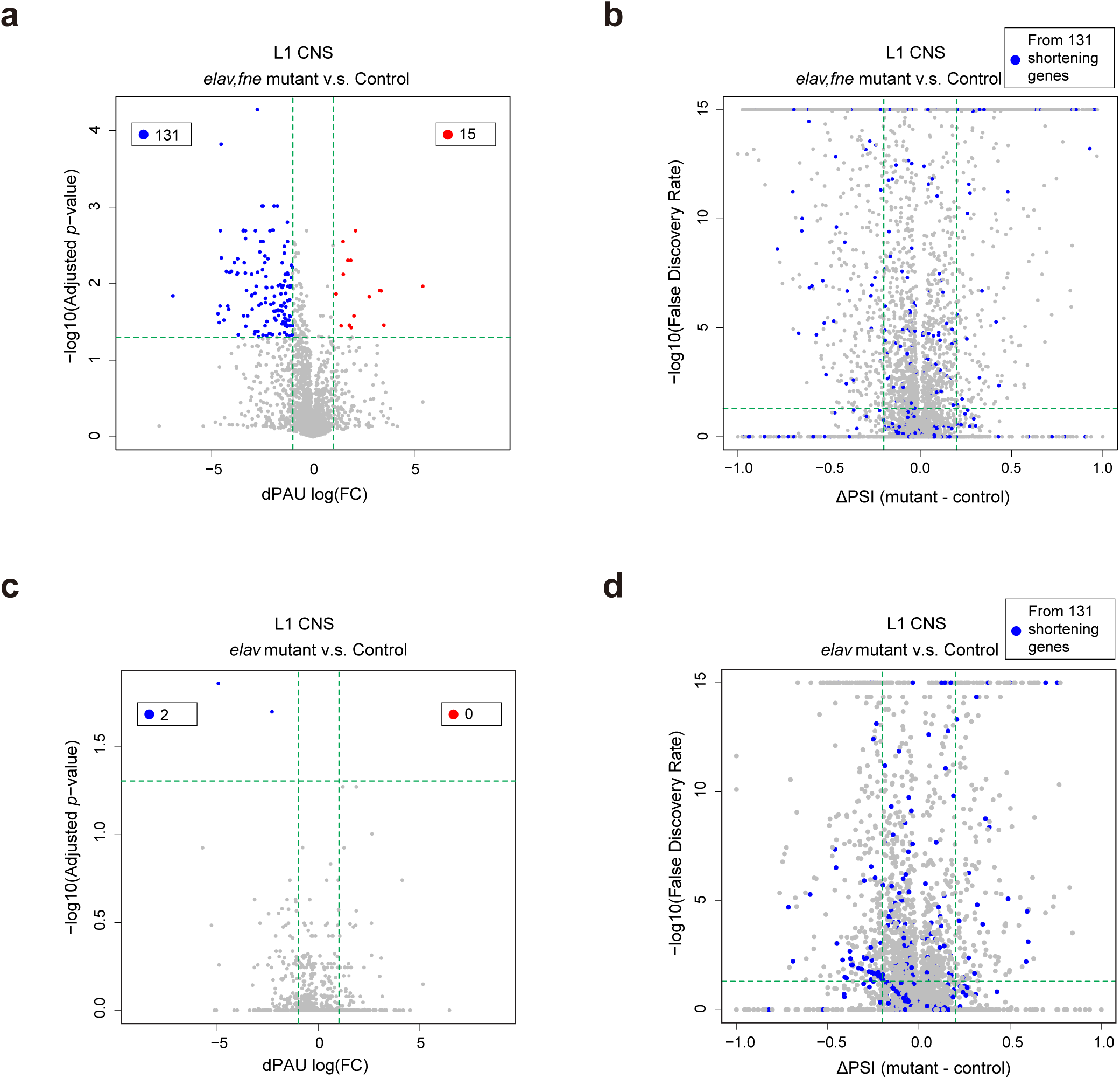
Short read RNA-Seq analysis of APA and CE AS in L1 CNS. (**a**) Short read RNA-Seq analysis of dPAU in L1 CNS samples from *elav,fne* mutant vs control using QAPA. 131 genes show significantly shortened 3’UTR in *elav,fne* mutant. (**b**) Change of CE events when *elav,fne* mutant is compared with control. The CE events from 131 3’UTR shortened genes from panel (**a**) are shown in blue. (**c**) Short read RNA-Seq analysis of dPAU in L1 CNS samples from *elav* mutant vs control using QAPA. 2 genes show significantly shortened 3’UTR in *elav* mutant. (**d**) Change of CE events when *elav* mutant is compared with control. The CE events from 131 3’UTR shortened genes from panel (**a**) are shown in blue. Horizontal dash lines in (**a**) and (**c**) indicate adjusted *p*=0.05, while in (**c**) and (**d**) indicate adjusted FDR=0.05. Veritical dash lines in (**a**) and (**c**) indicate FC=0.5 (left) and FC=2 (right), while in (**c**) and (**d**) indicate ΔPSI=-0.2 (left) and ΔPSI =0.2 (right).

## Supplementary Tables List

Supplementary Table S1: Primer sequences and coordinates (dm6).

Supplementary Table S2: CE exon number and coordinates for each AS-APA gene that was analyzed in this work.

Supplementary Table S3: Short-read RNA-Seq analysis summary for the 252 3’UTR lengthening genes and all regulated CE events from 58 out of 252 3’UTR lengthening genes in 2-4hr and 16-18hr embryos.

Supplementary Table S4: Short-read RNA-Seq analysis summary for the 290 3’UTR lengthening genes and all regulated CE events from 62 out of 290 3’UTR lengthening genes in adult ovaries and heads.

Supplementary Table S5: biotinylated probe sets for PL-Seq. 1 and 2, designed for AS-APA genes, Used in Fig. 2, 3, 4, and Supplementary Fig. 3, 4, and 5. Probe set 3 is used in Fig. 5 and 6.

Supplementary Table S6: Short-read RNA-Seq analysis summary for the 131 3’UTR lengthening genes and all regulated CE events from 29 out of 131 3’UTR lengthening genes in L1 CNS of *elav,fne* mutant and control samples.

Supplementary Table S7: Short-read RNA-Seq analysis summary for distal polyA site usage and all regulated CE events from 29 3’UTR lengthening genes (same as in Supplementary Table S6) in L1 CNS of *elav* mutant and control samples.

Supplementary Table S8: PL-Seq verified AS-APA genes and 7 of them have also reported to have coordinated AS and APA in previously short-read RNA-Seq data analysis of *elav,fne* mutant embryos (Carrasco et al., 2020).

Supplementary Table S9: PL-Seq data analysis results showing the read counts of CE splicing in and total (both inclusion and exclusion).

